# MetaMaps – Strain-level metagenomic assignment and compositional estimation for long reads

**DOI:** 10.1101/372474

**Authors:** Alexander Dilthey, Chirag Jain, Sergey Koren, Adam M. Phillippy

## Abstract

Metagenomic sequence classification should be fast, accurate and information-rich. Emerging long-read sequencing technologies promise to improve the balance between these factors but most existing methods were designed for short reads. MetaMaps is a new method, specifically developed for long reads, that combines the accuracy of slower alignment-based methods with the scalability of faster k-mer-based methods. Using an approximate mapping algorithm, it is capable of mapping a long-read metagenome to a comprehensive RefSeq database with >12,000 genomes in <30 GB or RAM on a laptop computer. Integrating these mappings with a probabilistic scoring scheme and EM-based estimation of sample composition, MetaMaps achieves >95% accuracy for species-level read assignment and *r*^2^ > 0.98 for the estimation of sample composition on both simulated and real data. Uniquely, MetaMaps outputs mapping locations and qualities for all classified reads, enabling functional studies (e.g. gene presence/absence) and the detection of novel species not present in the current database.

**Availability and Implementation:** MetaMaps is implemented in C++/Perl and freely available from https://github.com/DiltheyLab/MetaMaps (GPL v3).

## Introduction

Metagenomics, the study of microbial communities with the methods of genomics, has become an important tool for microbiology [1]. One key step in metagenomics is to determine the source genomes that a metagenomic sequencing dataset is derived from. This can be done either at the level of individual reads (read assignment or binning) or at the level of the complete dataset (compositional analysis).

A variety of methods have been developed for the analysis of metagenomic datasets, broadly falling into three classes. First, kmer-based read classifiers. This class includes approaches like Kraken [2], Opal [3], CLARK [4] and MetaOthello [5]. Second, alignment-based methods, for complete genomes or signature or marker genes. This category includes tools like Megan [6, 7], MetaPhlan [8], GASiC [9] and MG-RAST [10]. Third, Bayesian or EM-based estimators. This class includes Bracken [11], MetaKallisto [12] and Pathoscope [13, 14]. There are also approaches based on linear models or linear mixed models, for example PhyloPythia [15, 16], DiTASiC [17] and MetaPalette [18]; and methods that combine Markov models with alignment, for example Phymm/PhymmBL[19, 20]. The large majority of these methods have been designed for the analysis of short-read data; a small number of long-read-specific methods exist, but they are limited to specific data types (PacBio CCS [21]) or protein analysis (MEGAN-LR [22]).

The dominance of short-read sequencing in the field of metagenomics has traditionally been driven by cost efficiency. However, long-read sequencing (defined here as reads >2,000 bases) has recently become more cost-effective and has two intrinsic advantages over short-read sequencing for the interrogation of metagenomes. First, long reads preserve more long-range genomic information such as operon structures and gene-genome associations. The availability of this information can be key to functional and evolutionary studies, concerning, for example, the organization of metabolic pathways and horizontal gene transfer across metagenomes. Second, some long-read sequencers (the Oxford Nanopore MinION in particular) support rapid, portable and robust sequencing workflows, enabling “in-field” metagenomics. This is expanding the types of applications and scenarios that DNA sequencing and metagenomics can be applied to, such as the in-situ characterization of soil metagenomes in remote locations [23] or real-time pathogen sequencing during outbreaks [24]. For these reasons, the applicability and importance of long-read sequencing to metagenomics is growing rapidly.

This development of sequencing technology, however, has not yet been matched with the development of long-read-specific metagenomic analysis algorithms. Whereas tools that were designed for short reads can usually be applied to long-read sequencing datasets in principle, they often do not fully capitalize on the specific properties of the data. In short-read metagenomics, there are pronounced trade-offs between speed, accuracy and information richness. For example, methods like Kraken are very fast, but they do not attempt to determine the genomic positions of individual reads; alignment-based methods, on the other hand, can determine the genomic locations and alignment qualities of individual reads, but they are typically slow.

In the space of long-read metagenomics, desirable algorithms are both fast (to deal with large data volumes of incoming sequencing data on acceptable time scales, e.g. in the field) and produce highly informative output that includes per-read positional and quality information (because the availability of long-range spatial information is one of the key advantages of long-read sequencing). Here we show that this is indeed possible by leveraging the specific properties of long reads in a new approach called MetaMaps.

MetaMaps implements a two-stage analysis procedure. First, a list of possible mapping locations for each long read is generated using a minimizer-based approximate mapping strategy [25]. Second, each mapping location is scored probabilistically using a model developed here, and total sample composition is estimated using the EM algorithm. This step also enables the disambiguation of alternative read mapping locations.

Our approach has three main advantages. First, utilizing a mapping approach enables MetaMaps to determine individual read mapping locations, estimated alignment identities, and mapping qualities. These can be used, for example, to determine the presence of individual genes, or to assess the evidence for the presence of novel strains or species (which will exhibit systematically decreased alignment identities). Second, our approach is robust against the presence of large “contaminant” genomes, introduced during sample collection and processing or part of the environmental DNA, which often lead to false-positive classifications in methods that rely purely on individual k-mers. Third, reliance on approximate mapping makes the algorithm much faster than alignment-based methods, and our mapping algorithm can be tuned to different read lengths and qualities.

MetaMaps is also well-equipped to handle the continuous growth of reference database size [26]. First, MetaMaps implements a “limited memory” mode that, while leading to slightly increased runtimes, reduces memory usage while maintaining the same level of accuracy. This enables, for example, complete mapping of a long-read metagenomic sample to a comprehensive NCBI RefSeq database on a laptop computer. Second, by using the EM algorithm for borrowing information across reads [12, 14], MetaMaps can distinguish between closely related database genomes, a challenge that becomes more common as reference databases grow. The source package also includes support for Krona [27] and a set of lightweight R scripts for quick visualization of the sample-to-database mapping results.

## Materials and methods

### Strain-level metagenomics assignment

“Strain-level” accuracy is defined as source-genome resolution; that is, a read or a compositional estimate is counted as correct at strain-level resolution if and only if it is assigned to its correct genome of origin (known *a priori* for simulated data and determined via alignment for real data; see Methods). There are some instances in which a reference genome is directly attached to a species node in the NCBI taxonomy (typically when there is only one reference genome for the species); for these genomes, strain- and species-level resolution are synonymous.

### Reference database and strain-specific taxonomy

A comprehensive reference database, comprising 12,058 complete RefSeq genomes and 25 gigabases of sequence, is used for all experiments presented here. It includes 215 archaeal, 5,774 bacterial, 6,059 viral/viroidal, and 10 eukaryotic genomes, 7 of which are fungal and one of which is the human genome. The database also includes an extended, strain-specific version of the NCBI taxonomy, in which each input genome maps to exactly one node (see Supplementary Figure S1). MetaMaps supports the use of custom databases, and scripts for downloading and processing genomes from NCBI are part of the distribution.

### Initial read mapping and identity estimation

MetaMaps employs a fast approximate long-read mapping algorithm to generate an initial set of read mappings, fully described elsewhere [25] and visualized in Supplementary Figure S2. Briefly, a minimizer [28] index is constructed from the reference database; intersections between the minimizer sets selected from a sequencing read and the reference define an initial set of candidate mapping locations. Low-identity candidate mapping locations are eliminated using a fast linear-time algorithm. For all surviving *N* candidate mapping locations of a read *r*, alignment identity is estimated using a winnowed-minhash statistical model [25]. Briefly, let *S_r_* be the MinHash sketch [29] of the read-selected minimizers; let *S_i_* be the MinHash sketch the set of reference-selected minimizers for mapping location 1 ≤ *i* ≤; let *S_r_*_**∪***i*_ be the MinHash sketch of the minimizer set union. We define 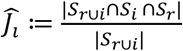 as an estimate of the Jaccard similarity between the k-mer sets of the mapping location *i* and the read, and further 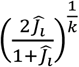 as an estimator of the corresponding alignment identity for k-mer size *k*. Minimizer density is auto-tuned [25] based on a user-defined minimum read length and minimum mapping identity (by default and for all experiments presented here: 2,000 bases and 80% identity).

### Mapping qualities

Using the MinHash sketch, we define a probabilistic mapping quality model to quantify mapping uncertainty for reads with multiple mapping locations. Under the assumptions of the model and conditional on a known sequencing error rate, we model P(|*S_r_*_**∪***i*_ ∩ *S_i_* ∩ *S_r_* |) as binomial with parameters *n* = |*S*_*r*∪*i*_| and k-mer survival rate P = (1 − *e*)^*k*^. The posterior probability (mapping quality) of mapping location for read *r* is defined as 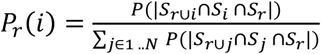 *e* is unknown and read-specific; for simplicity we define (1 − *e*) as the estimated identity of the highest-scoring mapping for each read.

### Sample composition and read redistribution

Let *G* be the set of genomes in the database and let *F_g_* be the (unknown) probability that a sequencing read in the sample emanates from database genome *g* **∈** *G*. *F* is a vector that describes the sample composition and is to be estimated.

We define the likelihood of the mapped read set *R* as 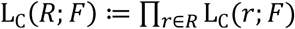 and the likelihood of an individual read *r* as

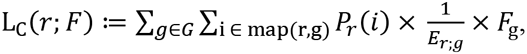
where *i* is a read mapping location. To account for genomic duplications, map(r,g) is the set of all mapping locations of read *r* in genome *g*, *P_r_*(*i*) is the posterior probability of mapping location *i*, and *E_r;g_* is the count of effective start positions for read *r* in genome *g*. *E_r_*_;*i*_ is defined as contig length minus read length, summed over all contigs of genome *g*. This implies a uniform distribution over possible within-genome start positions of reads; for simplicity, we don’t distinguish between circular and non-circular contigs.

We define the composition-dependent posterior probability of mapping location *i* as

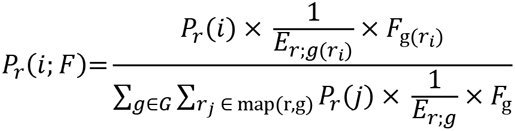
where g(*r_i_*) is the genome associated with mapping location *i*. Summing over all reads, we obtain an updated sample composition estimate as

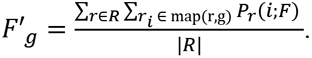

The frequency vector *F* is initialized with 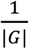 for each element and we update until convergence of the likelihood L_C_(*R*; *F*). Unmapped reads exceeding the minimum read length are assigned to the ‘unassigned’ category (special taxon ID 0), followed by a final renormalization. Each mapped read *r* is individually assigned to its maximum likelihood genome location.

### Memory-efficient mapping

To enable classification against large reference databases with limited resources, MetaMaps supports the specification of a maximum memory target amount (“memory-efficient mapping”). When run in memory-efficient mode, the order of contigs in the reference database is randomized and processed in a sequential manner. Starting from the first contig, index construction is started and continues until internally estimated memory consumption is just below the user-specified target amount or until the end of the reference database has been reached. The input data are then mapped against this index representing a subset of the reference database and stored on disk. The index is cleared, and construction of a new index begins at the position at which the process was previously aborted. Suboptimal mapping locations will later be assigned low mapping qualities during the EM step.

### Kraken and Bracken

We compare MetaMaps to Kraken [2] and Bracken [11], two archetypical tools for taxonomic assignment of reads (Kraken) and for sample composition estimation (Bracken). For each experiment, we use the same database for Kraken, Bracken (with `--read-length=2000`) and MetaMaps. Kraken returns an explicit taxonomic assignment for each read and returns the special taxon ID 0 if a read remains unclassified. Bracken uses a Bayesian model to update and refine the read counts at lower levels of the taxonomy, so we call Bracken separately for re-estimation at the levels of species, genus, and family, and obtain a sample composition estimate by dividing the number of assigned reads per node by the number of total reads in the input dataset. After summing over all nodes, remaining reads are counted towads taxon ID 0.

### Evaluation experiments

We carry out four experiments to evaluate MetaMaps. For read simulation, we use pbsim [30] with parameters `--data-type CLR --length-mean 5000 --accuracy-mean 0.88`.

#### Experiment i100

The basis for this experiment is the “medium complexity” metagenome scenario described in [31], specifying a metagenome of 100 species with defined frequencies. The most recent records for the specified sequence accession numbers, all of which are present in the MetaMaps database, are obtained via the NCBI query interface, omitting 4 that were removed from RefSeq since publication of [31]. We use pbsim with the above parameters to simulate 1 gigabase of long-read data from the 96 genomes. The utilized accessions and the realized read counts are shown in Supplementary Table S3.

#### Experiment p25

The basis for this experiment are 25 bacterial genomes [13] comprising 5 strains of *Escherichia coli*, 5 strains of *Staphylococcus aureus*, 5 strains of *Streptococcus pneumoniae* and 10 other common bacterial strains, all of which are present in the MetaMaps reference database. We arbitrarily assign genome abundances according to a log-normal model and use pbsim to simulate 1 gigabase of long-read sequencing data. The utilized accessions and the realized read counts are listed in Supplementary Table S4.

#### Experiment e2

As a negative control, we use pbsim (with additional parameter --length-min 2100 to ensure mappability of all simulated reads) to simulate long-read sequencing data from two eukaryotic genomes not present in the MetaMaps or Kraken databases. Specifically, we simulate 1 gigabase of sequencing data from the *Aedes aegypti* (yellow fever mosquito) genome (GCF_002204515.2), and 1 gigabase of sequencing data from the *Toxoplasma gondii* ME49 genome (GCF_000006565.2). The two read sets are analyzed independently with MetaMaps and Kraken/Bracken.

#### Experiment HMP7

For the HMP7 experiment, we use data from a single flow cell (2.9 Gb FASTQ; 319,703 reads) of the PacBio HMP sequencing experiment (Set 7), based on genomic DNA (Even, High Concentration) from Microbial Mock Community B. To generate a truth set, we use bwa mem [32] with `-x pacbio` to map the reads against the reference genomes specified for the DNA source. All reads that cannot be mapped with bwa are excluded, and the primary alignment for each read determines the assumed true placement. The accession numbers of the reference genomes and their realized read counts are listed in Supplementary Table S5.

#### Evaluation metrics

Read assignment accuracy is assessed by PPV (the proportion of correct read assignments) and recall (the proportion of reads having been given a correct assignment). Specifically, there is a true taxonomic assignment *truth*(*r*) **∈** {0 **∪** T}for all reads *r* in the validation set and an inferred taxonomic assignment *inference* (*r*) ∈ {0 ∪ *T*} for some or all reads *r* in the validation set, where *T* is the set of nodes in the database taxonomy and 0 is a special taxon ID indicating unassignedness. The function *to_level*(*a*,*l*), where *a* ∈ {0 ∪ *T*} is a taxonomic assignment and *l* a taxonomic level, enables the conversion of these assignments to specific taxonomic levels and is defined as

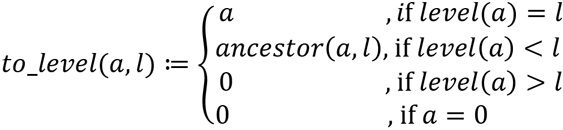
where *level*(*a*) is a function that returns the taxonomic level of an assignment *a* and where (*a*, *l*) is a function that returns the *l*-level ancestral node of *a*. An assignment is counted as correct at level *l* iff *to*_*level*(*inference*(*r*), *l*) = *to*_*level*(*truth*(*r*), *l*). For MetaMaps, strain-level correctness is defined as *inference*(*r*)= *truth*(*r*). PPV at level *l* is defined as the proportion of correct assignments at level *l*, and recall at level *l* is defined as the number of correct assignments at level *l* divided by the total number of reads in the validation set. MetaMaps produces assignments for all reads in the validation set longer than 2000bp, some of which might be 0 (see above). Kraken produces assignments for all reads in the validation set, some of which might be 0 or, because they correspond to non-leaf nodes of the taxonomy, will be converted to 0 when evaluating at lower taxonomic levels.

Note that the presence of 0s is not confined to the inference set; *to*_*level*(*truth*(*r*), *l*) is 0 if the taxonomic level of *truth*(*r*) is higher than *l*. This happens in experiments e2 and HMP7 for reads that emanate from out-of-database genomes; for these, *truth*(*r*) is set to the node that represents the most recent common ancestor between the source and database genomes (see below and Supplementary Figure S6). Under the model of Kraken, reads assigned to higher nodes in the taxonomy could also be considered as entirely uncalled at lower levels; we therefore also report metric PPV2, which, for level *l*, is defined as the proportion of correct non-0 assignments at level *l*.

Sample composition estimation accuracy is assessed by the two metrics L1 (the distance between the true and inferred sample composition vectors using the L1 norm) and r2 (Pearson’s *r*^2^ for the true and inferred composition vectors); both composition metrics are computed over columns that are non-0 in either vector.

#### Evaluating HMP7 performance

HMP7 comprises 20 microbial strains. Of note, 3 of these are not part of the MetaMaps database, because the corresponding assembly records as specified in the Mock Community B Product Information Sheet have been removed from RefSeq or are not classified as “complete genome”. For two missing genomes (*Acinetobacter baumannii ATCC 17978* and *Rhodobacter sphaeroides 2.4.1*), the MetaMaps database contains closely related genomes of the same species (mash distances [33] 0.00 and 0.01, respectively). For the remaining genome (*Actinomyces odontolyticus ATCC 17982*), the next-closest in-database relative belongs to a different species (*Actinomyces meyeri*, mash distance 0.14). For the truth set that read assignments are evaluated against, out-of-database genomes are assigned to non-leaf nodes of the taxonomy (see above); specifically, the true strain-level taxon IDs for all *Acinetobacter* and *Rhodobacter* reads, and the true strain- and species-level taxon IDs for all *Actinomyces* reads, are set to 0 (Supplementary Figure S6). Note that MetaMaps will always assign all mapped reads at the strain level, whereas Kraken can place reads at higher taxonomic levels.

## Results

### Simulated data

We first evaluate the performance of MetaMaps in two simulation experiments. Experiment i100 represents a “medium-complexity” metagenomic analysis scenario with approximately 100 species; experiment p25 a “pathogenic” metagenomic scenario with 15 strains of 5 potentially pathogenic bacteria and 10 other bacterial strains. At the strain level, MetaMaps assignments achieve a PPV of 92% (i100) and 86% (p25). At the species level, PPV increases to 98% (i100) and 100% (p25). MetaMaps consistently outperforms Kraken in terms of PPV by a margin of 4 – 10% (species, genus); at the family level, MetaMaps still outperforms Kraken, but by <1%. MetaMaps also consistently outperforms Kraken in terms of PPV2, apart from the species level in i100, where its PPV2 is 0.2% lower than that of Kraken (though at a 6% higher recall). Recall for MetaMaps is consistently about 3% lower than PPV, driven by the minimum length requirement. At the species level, MetaMaps outperforms Kraken by 1 – 6% in terms of recall; at higher taxonomic levels, Kraken recall is 1 – 3% above that of MetaMaps. For reads longer than 2000 bp, MetaMaps achieves a recall of 99.9% or higher from the genus (i100) and species (p25) level onwards, consistently higher than that of Kraken. Read classification results are visualized in Figure 1 and full results are given in Supplementary Table S7.

**Figure 1.**
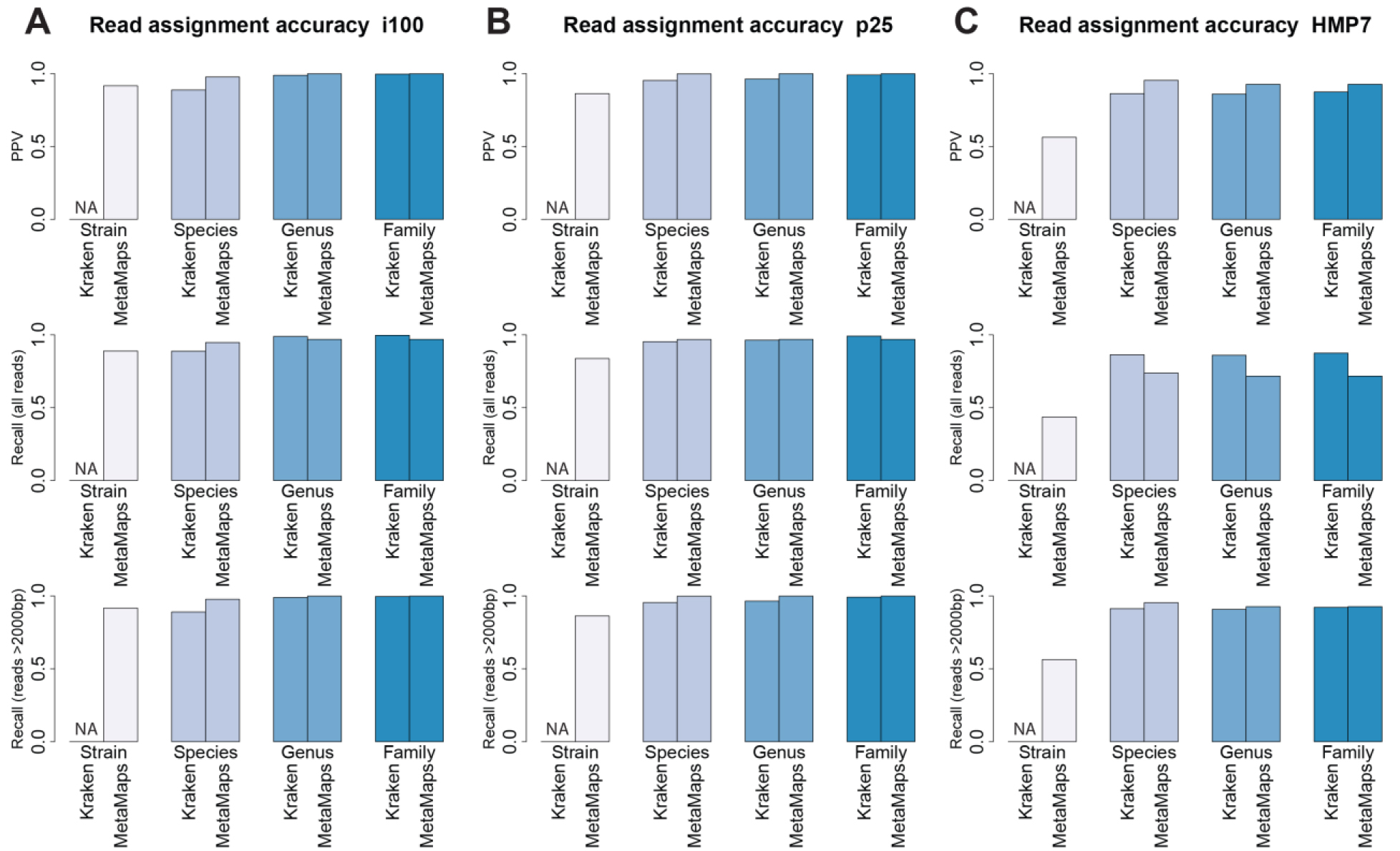
Read assignment accuracy in experiments i100 (panel A), p25 (panel B), and HMP7 (panel C) for Kraken and MetaMaps. Bar plots show PPV, recall for all reads, and recall for reads longer than 2000 bases at different evaluation levels. Note that Kraken was not designed to achieve strain-level resolution; it is therefore not validated at this level.

MetaMaps can also accurately estimate sample composition (Figure 2). At the strain level, MetaMaps achieves a Pearson’s *r*^2^ between estimated and true abundances of 0.87 (i100) and 0.77 (p25). These increase to >0.99 at higher levels. The performance of Bracken and MetaMaps is similar; MetaMaps, however, exhibits slightly smaller distances (L1-norm) between estimated and true compositions. Full compositional estimation accuracy results are shown in Supplementary Table S8.

**Figure 2.**
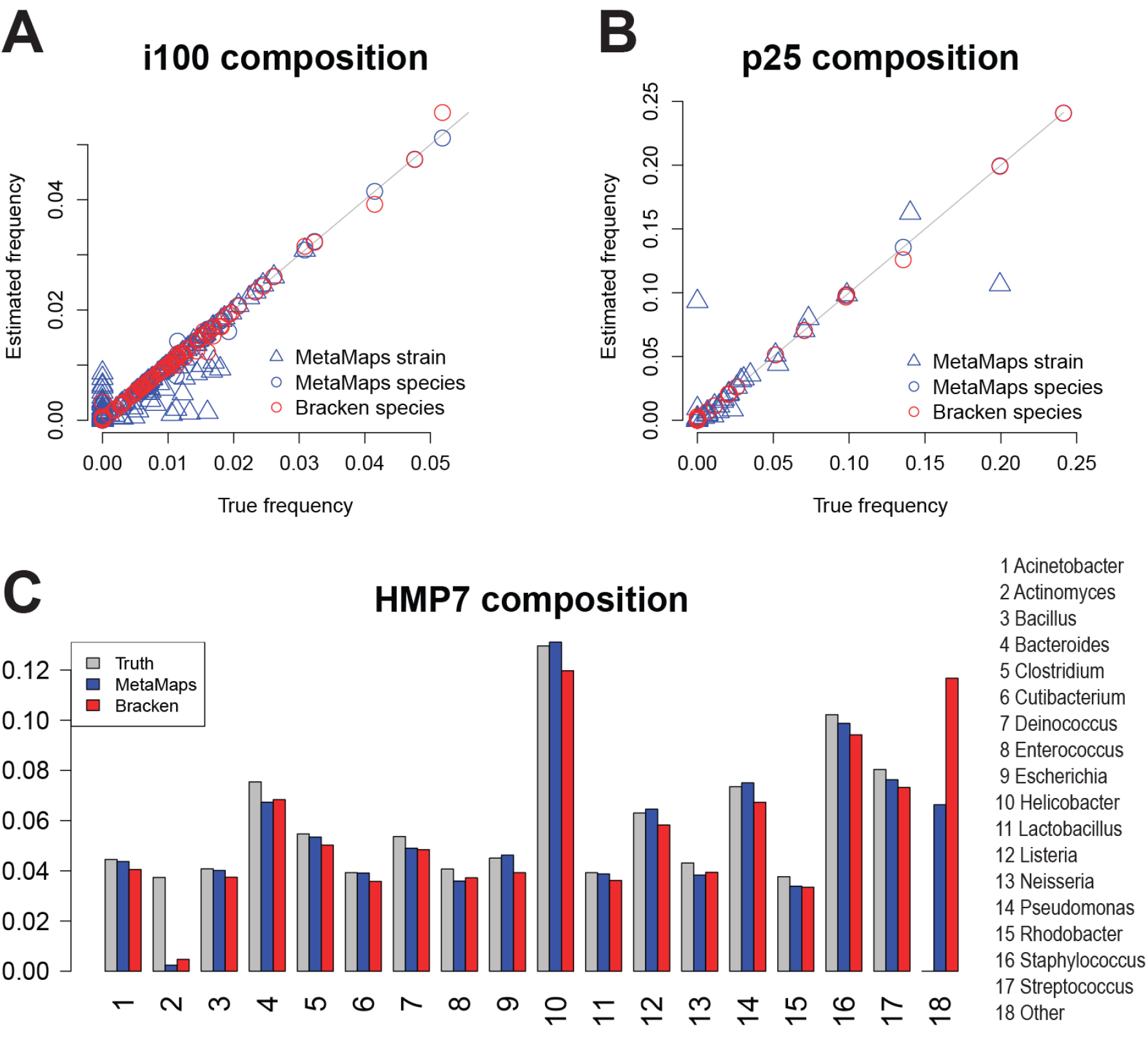
Panels A and B: True and inferred sample composition frequencies, for MetaMaps and Bracken. Bracken was not designed to achieve strain-level accuracy and is therefore not validated at this level. Panel C: Compositional estimation in the HMP7 experiment at the genus level, shown for Bracken and MetaMaps, and compared to the assumed true composition. Note that the *Acinetobacter*, *Actinomyces*, and *Rhodobacter* genomes present in the sequencing sample are not part of the database; reads classified as belonging to these genera map to other genomes of the same species / genus (see “Evaluating HMP7” in “Methods”).

We use the i100 experiment to assess the effect of read length on the ability to accurately classify a read. All methods show a trend towards higher classification accuracy for longer reads. This effect is most pronounced for Kraken (Figure 3), whereas MetaMaps exhibits relatively constant classification accuracy, once the minimum read length has been reached (see Discussion).

**Figure 3.**
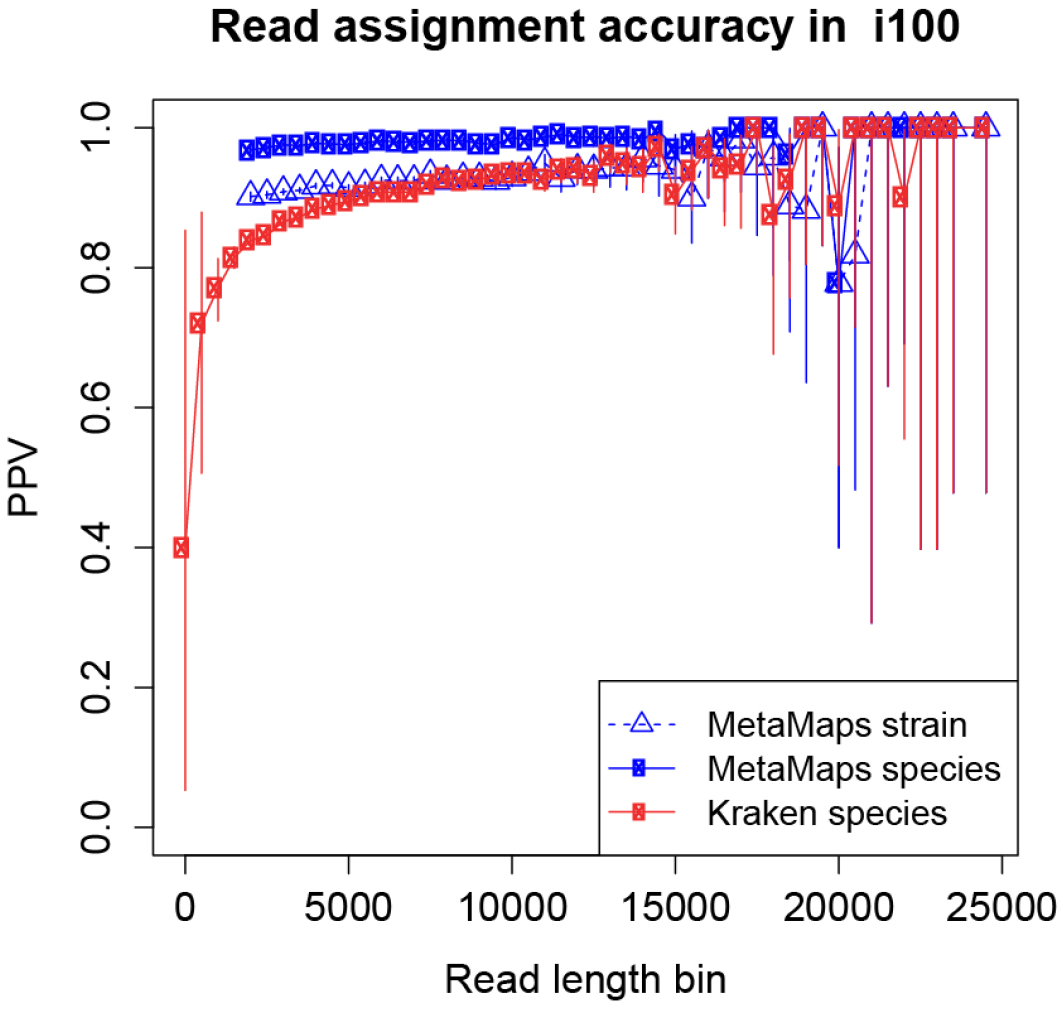
PPV of called reads in simulation experiment i100, stratified by read length. Note that MetaMaps results start at a minimum read length of 2,000, corresponding to the “minimum read length” parameter the algorithm was run with. For bins above the MetaMaps minimum read length, recall equals PPV.

### Real HMP7 data

To evaluate performance on real data, we apply MetaMaps and Kraken/Bracken to PacBio data from the Microbial Mock Community B of the Human Microbiome Project (HMP Set 7). Note that not all genomes represented in HMP7 are part of the utilized reference database (see Methods), and that sample-database mismatches are a recurrent concern in metagenomics.

First, we evaluate read assignment accuracy (Figure 1 and Supplementary Table S7). At the strain level, MetaMaps achieves a PPV of 56%; but we note that strain-level differences might exist between the sequenced HMP7 sample and the reference genomes deposited in NCBI. Consistent with this, PPV increases to 95% at the species level. MetaMaps consistently outperforms Kraken in terms of PPV, by a margin between 9% (species) and 5% (family). It also outperforms Kraken in terms of PPV2, though by smaller margins (1 – 2%). In the HMP7 dataset, a higher proportion (23%) of reads remain unassigned due to the minimum length requirement of MetaMaps, and Kraken achieves higher recall values than MetaMaps (margin 13% - 16%). We note, however, that MetaMaps recall for reads longer than 2000bp is >92% across all evaluated levels and consistently higher than that of Kraken (on the same set of reads).

Second, we consider the accuracy of sample composition estimation (Supplementary Table S8). As before, estimating sample composition at the strain level is most challenging (MetaMaps *r*^2^ = 0.3); the accuracy of the estimation is much higher at the species (*r* ^2^ = 0.98) and genus/family (*r* ^2^ = 0.91) levels. On the HMP7 data, MetaMaps has a consistent advantage over Bracken (Figure 3), which has a species-level *r*^2^ of 0.85. Of note, accuracy for the *Actinomyces* genus is low for both MetaMaps and Kraken, because the specific strain present in HMP7 is not part of the reference database (see Methods and next section).

### Database-sample mismatches

Mismatches between the sequencing sample and the utilized database are an important concern in metagenomics (i.e. sequencing reads originating from genomes not in the database). To evaluate the effect of large out-of-database genomes, we assess classification accuracy in experiment e2. Experiment e2 contains simulated reads from two eukaryotic genomes, neither of which is present in the reference database (the yellow fever mosquito and *Toxoplasma gondii*, representing plausible contamination scenarios; see Methods). For both read sets, MetaMaps has a low false-positive rate and correctly leaves the large majority of reads unclassified (99% PPV/recall at the species, genus and family levels for mosquito reads and 98% of toxoplasma reads); of note, the minimum length requirement of MetaMaps does not contribute to this result, as all simulated reads in this experiment are long enough (see Methods). Kraken and Bracken, on the other hand, falsely classify four times as many mosquito reads and eightfold more toxoplasma reads than MetaMaps (96% PPV/recall for mosquito reads and 83% for toxoplasma reads, at the same levels). Actual microbial contamination of these eukaryotic assemblies is possible, but given that MetaMaps shows similar sensitivity to Kraken on the other datasets, it is likely that the majority of Kraken/Bracken e2 calls are false.

We use the HMP7 data to evaluate the effect of subtler mismatches and whether the availability of read mapping locations and estimated alignment identities enable the detection of database-sample mismatches. MetaMaps provides an R tool to visualize spatial coverage and identities of the read mappings. Examination of these plots for the *Actinomyces* genome (Figure 4), which is diverged from the strain in the sequencing dataset (mash distance 0.14, see Methods), reveals both a highly uneven coverage pattern as well as a stark shift of read identities away from the expected average of around 0.88 (approximately equal to 1 minus the sequencing error rate, Supplementary File S9). It is clear from these results that any *Actinomyces*-related result from this experiment would have to be interpreted with caution, consistent with the high evolutionary distance between the sample *Actinomyces* genome and its next-closest relative in the database.

**Figure 4.**
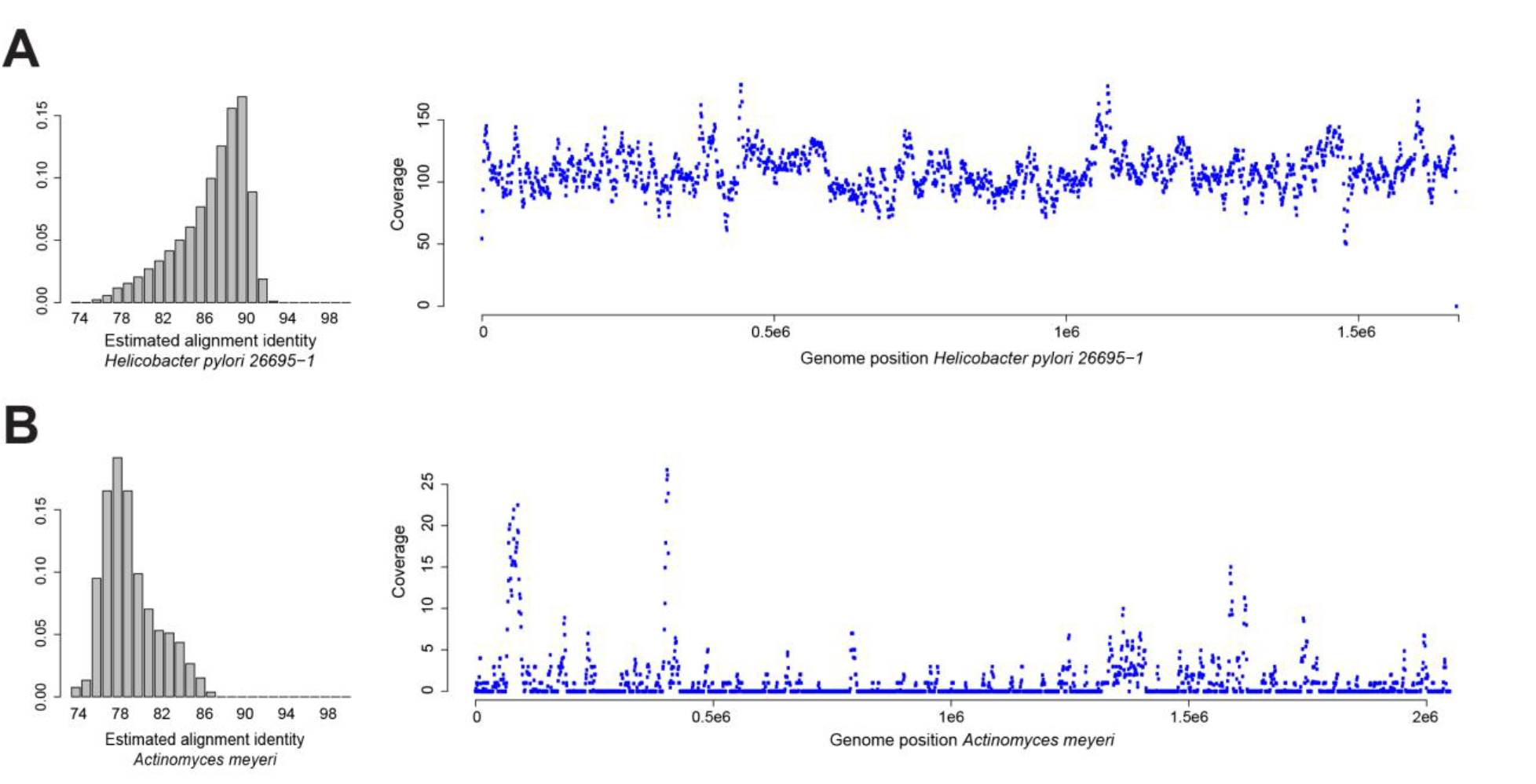
Estimated alignment identity and spatial genome coverage for *Helicobacter pylori 26695-1* (panel A) and *Actinomyces meyeri* (panel B) in the HMP7 experiment. *Actinomyces meyeri* is the next-closest database relative for the *Actinomyces odontolyticus* genome present in the HMP7 sequencing data (mash distance 0.14). *Helicobacter pylori 26695-1* is present in HMP7 and in the reference database. Uneven spatial coverage and estimated mapping identities shifted away from the expected mode around 0.88 for *Actinomyces meyeri* are indicative of a mismatch between the sequencing sample and the reference database. Complete plots for HMP7 are contained in Supplementary File S9.

### Runtime and memory-efficient mode

At the expense of slightly increased runtimes, MetaMaps can be run in memory-efficient mode, with an upper memory consumption target to be specified by the user. We evaluate this mode by repeating the i100 experiment with target memory set to 20 GB. First, the complete MetaMaps classification process in standard mode takes 9 CPU hours and memory consumption peaks at 139 GB, well above the capacity of standard workstation computers. With a target memory set to 20 GB, the classification process takes 15 CPU hours and memory peaks at 26 GB, a requirement that can be satisfied by medium-ranged workstations (Supplementary Table S10). Note that effectively consumed memory can exceed the specified target maximum amount (Methods). The accuracy of both read assignment and sample composition is virtually unaffected by limiting memory (Supplementary Table S7 and Supplementary Table S8). Kraken/Bracken are 1–2 orders of magnitude faster than MetaMaps (Supplementary Table S10) but require more memory (154 GB) and do not report read mapping information.

## Discussion

We have presented MetaMaps, an algorithm specifically developed for the analysis of long-read metagenomic datasets that enables simultaneous read assignment and sample composition analysis. The key novelty of MetaMaps is the combination of an approximate mapping algorithm with a model for mapping qualities and the application of the EM algorithm for estimation of overall sample composition. As discussed in the Introduction, this design was motivated by the aim to develop an algorithm tailored for long reads that is both fast and preserves per-read spatial and quality information.

Our evaluations show that MetaMaps outperforms Bracken in terms of sample composition estimation, that the read assignments of MetaMaps are more accurate than those of Kraken, and that recall for long reads (above defined as > 2000 bp) is consistently higher than that of Kraken. However, a proportion of reads remain unassigned under the MetaMaps model because they do not meet the minimum length requirement. This is a direct consequence of the approach we chose for approximate mapping, which determines minimizer density based on expected read lengths and alignment identities. Reads that fall below the chosen minimum length end up with minimizer sets that are too small to reliably determine their mapping locations. It is worth noting, however, that minimum read length and expected alignment identities are user-defined parameters that can be set empirically (for example based on the distribution of read lengths) and according to user preferences (e.g. with respect to runtime and the proportion of reads that remain unclassified). In addition, read lengths can be optimized with specific protocols for the extraction of high-molecular-weight DNA; the applicability of these, however, depends on sample and experimental conditions.

MetaMaps computes a maximum likelihood approximate mapping location, an estimated identity and mapping qualities for all candidate mapping locations. Its output is nearly as rich as alignment-based methods and enables a very similar set of applications, while being many times faster.

We have demonstrated the advantages of this approach. First, MetaMaps is robust against the presence of large out-of-database genomes, for example eukaryotic genomes. Contamination and environmental DNA are important concerns in many metagenomic studies, and the MetaMaps model is more robust against these than methods based purely on individual k-mers. Second, estimated alignment identities can be informative about the presence of novel species or strains that have a related in-database genome. This too is an important concern, as microbial reference databases comprise but a fraction of total microbial genome diversity. Third, because it reports mapping information, MetaMaps can be used to ascertain the presence of particular genes or loci of interest, for example antibiotic resistance genes or virulence factors.

In many plausible metagenomic analysis scenarios, computing resources are limited — for example when sequencing metagenomes using a portable nanopore device during a field trip without reliable internet connection. We therefore developed a feature to limit memory consumption during the approximate mapping step. As we showed, reducing memory consumption comes at a runtime cost, but accuracy remains unaffected, and, in contrast to many other similar approaches, classification is still carried out against the complete reference database.

There are two important directions for future work. First, building support for streaming data into MetaMaps would be an important feature for many clinical applications. It could also be used to control the “read until” {Loose, 2016 #190} feature of the Nanopore technology. Such an extension would be relatively straightforward to implement in terms of the algorithms’ architecture by dynamically re-computing mapping qualities, to the extent that they are influenced by changes in the global sample composition frequency vector. Second, it would be desirable to integrate an explicit term for genomic divergence directly into the statistical models of MetaMaps; this would enable the explicit detection of and testing for novel strains and species in the sequencing sample. K-mer painting approaches [18] have been suggested as a solution to this problem in the short-read space. How to best implement the detection of novelty from long-read data remains an open question for further research.

## Supplementary Figure Legends

**Supplementary Figure S1.** MetaMaps uses an extended version of the NCBI taxonomy in which each reference database genome has a unique taxon ID. This is constructed by creating additional pseudo taxon IDs (prefixed with an ‘x’), which distinguish between genomes attached to the same node in the original NCBI taxonomy.

**Supplementary Figure S2.** High-level overview of the approximate mapping algorithm [25]. Minimizers are selected from the reference and from the reads. Minimizer matches between read and reference are identified using a hash table, inducing candidate mapping locations. Minimizer density is determined based on minimum read length and alignment identity. For each candidate mapping location, we use a winnowed-minhash approach, based on read and reference minimizers, to estimate the Jaccard similarity between the full kmer sets of the read and the candidate mapping location, and convert this estimate into an estimate of alignment identity. The steps below the dashed line show the subsequent steps of mapping quality computation and EM-based sample composition estimation.

**Supplementary Table S3.** Summary of the i100 simulated data, representing a medium-complexity metagenome of 96 species (see Methods). “taxonID” and “Name” specify the NCBI taxonomy ID and the name of the organism, “NCs” the contig IDs that were used for read simulation (with pbsim; see Methods). “Bases”, “nReads” and “Genomes” refer to the number of simulated bases, reads and genome equivalents per organism.

**Supplementary Table S4.** Summary of the p25 simulated data, representing 15 potentially pathogenic and 10 common bacterial species (see Methods). “taxonID” and “Name” specify the NCBI taxonomy ID and the name of the organism, “NCs” the contig IDs that were used for read simulation (with pbsim; see Methods). “Bases”, “nReads” and “Genomes” refer to the number of simulated bases, reads and genome equivalents per organism.

**Supplementary Table S5.** Summary of the HMP7 data, which were generated by sequencing a mock community sample generated by the Human Microbiome Project with the PacBio technology. To generate a truth set, all reads were mapped using bwa against the reference genomes specified in the sample product information sheet (column “GIs used for truth-set mapping”). “taxonID” and “Name” specify the NCBI taxonomy ID and name of the organism; “Bases” and “nReads” specify, for each organism, the sum of read lengths and the absolute read count assigned to each organism in the truth set; “Genomes” is the base count divided by approximate genome length.

**Supplementary Figure S6.** In experiment HMP7, not all genomes present in the input data are present in the reference database. We assign reads that emanate from out-of-database entities to the taxonomic node that represents the most recent common ancestor of the read’s source genome and its next-closest database relative; and for all taxonomic levels below the most-recent-common-ancestor node, true read assignment is defined as “Unassigned” (special taxon ID 0).

**Supplementary Table S7.** Read assignment accuracy at different evaluation levels for the p25, i100 and HMP7 experiments. Strain-level accuracy is measured at the level of individual database genomes (see Methods). Kraken doesn’t support strain-level analysis and is therefore not validated at this level. “# Reads” specifies the total size of the input read set; “PPV (full)” is the proportion of correct read assignments; “PPV2 (!= 0)” is the proportion of correct non-0 read assignments; “Recall” is the proportion of reads having received a correct call.

**Supplementary Table S8.** Accuracy of compositional estimation at different evaluation levels for the p25, i100, and HMP7 experiments. Strain-level accuracy is measured at the level of individual database genomes (see Methods). Bracken was not designed to achieve strain-level resolution and Kraken was not designed for compositional estimation; the corresponding cells are therefore grayed out. Metric L1 quantifies the difference between the true and inferred composition vectors using the L1 norm; metric r2 quantifies the similarity between the true and inferred composition vectors using Pearson’s *r*^2^; both metrics are limited to columns that are non-0 in either the true or the inferred composition vector.

**Supplementary File S9.** Summary statistics (read length, estimated alignment identities, and genome coverage for each genome with an estimated frequency >0.1%; alternating colors in the coverage plots indicate chromosome boundaries) for the HMP7 analysis. Note how the *Actinomyces* genome differs from the other genomes in terms of alignment identities and spatial genome coverage. MetaMaps comes with a lightweight R script for the generation of equivalent plots for user datasets.

**Supplementary Table S10.** CPU time and peak memory statistics for the p25, i100 and HMP7 experiments. The i100 “limited memory” experiment was run with a target maximum memory amount of 20GB (`--maxmemory 20`).

